# Can the metacommunity data matrix predict changes in species incidence and abundance?

**DOI:** 10.1101/047696

**Authors:** Donald M. Waller, Erika L. Mudrak, David A. Rogers

**Author notes:** Current Address: Cornell University Statistical Consulting Unit, Ithaca, NY, USA.

## Abstract

Metacommunity matrices contain data on species incidence or abundance across sites, compactly portraying community composition and how it varies over sites. We constructed models based on an initial metacommunity matrix of either species incidence or abundance to test whether such data suffice to predict subsequent changes in incidence or abundance at each site. We then tested these models against extensive empirical data on vascular plant incidence and abundance collected from 156 forested sites in both the 1950s and 2000s. Predictions from these models parallel observed changes in species incidence and abundance in two distinctly different forest metacommunities and differ greatly from null model predictions. The abundance model shows greater power than the incidence model reflecting its higher information content. Predictions were more accurate for the more diverse forests of southern Wisconsin which are changing faster in response to succession and fragmentation. Simulations demonstrate that these results are fairly robust to variation in sampling intensity. These models, based only on the metacommunity matrix, do not require data on site conditions or species' characteristics. They thus provide a useful baseline for assessing more complex models incorporating data on species' functional traits, local site conditions, or landscape context. They may also prove useful to conservation biologists seeking to predict local population declines and extinction risks.

## Introduction

Ecologists seek to understand community assembly in terms of external forces (e.g., disturbance, edaphic factors, etc.), adaptations to local environmental conditions ("species sorting"), competitive niche-based processes, trophic interactions, and stochastic factors. Shmida and Wilson (1985) first sought to identify the overall mechanisms affecting community structure and diversity. Their first mechanism reflects differences among species in local resource use, or niche differences. Their second concerns environmental differences among sites in abiotic habitat conditions. Traditional plant ecology has focused on these two, but Shmida and Wilson also noted two additional mechanisms reflecting the stochastic processes that occur within and among sites. When sites are connected via dispersing individuals, within-site species dynamics are affected by "mass effects" related to regional site occupancy and abundance. They defined mass effects as occurring when species establish in sites where they cannot maintain themselves and when individuals flow from areas of high success to less favorable areas. These ideas also emerged in the "rescue effect" of Brown and Kodric-Brown (1977). They further noted that even ecologically equivalent species could stably coexist under certain conditions, anticipating the neutral models of Bell (1991) and Hubbell (1997, 2001). Those authors developed these ideas to explore how neutral models based on ecologically equivalency can be used to predict community structure and diversity. These predictions match patterns observed in many plant and animal communities (Chave et al. 2002, Hubbell 2006) despite limited empirical support for certain key assumptions (McGill et al. 2006). This may reflect the fact that ecological mechanisms often operate to reduce species to similar fitness levels (Chave 2004).

Metacommunities represent an ensemble of the communities similar enough to share a common species pool and close enough to influence each other's composition via dispersal and colonization (Wilson 1992). In general, we expect patch heterogeneity and dispersal to influence local patterns of occupancy and abundance. Such effects have been observed in experimental microcosms (Davies et al. 2009) and field plant populations (Turnbull et al. 2000) but not populations of tree hole mosquitoes (Ellis et al. 2006). In reviewing bird and reptile distributions, Driscoll and Lindenmayer (1979) found no consistent metacommunity response. Logue et al. (2011) further concluded that "At present, [our] understanding of metacommunity dynamics is predominantly theoretical in nature."

Leibold et al. (2004) also identified four major metacommunity paradigms that included mass effects, neutral models, species interactions (within-site dynamics driven by niche-based competitive interactions – e.g., Gravel et al. 2006), and patch-dynamics driven by dispersal-competition trade-offs among species. They and Leibold and Miller (2004) explicitly associate mass effects with high dispersal and heterogeneous habitat conditions. Having a high regional abundance can lead to both greater occupancy, as empty sites are more quickly colonized, and unsuitable sites may support sink populations. Heterogeneity in site conditions is invoked as the mechanism driving variation in species abundances. Variable site conditions, in turn, drive individuals to emigrate from densely occupied sites to unoccupied or low density sites. Note, however, that stochastic colonization, population growth, and random disturbances also drive variation in species abundances even among homogenous sites. Thus, we need not invoke habitat differences among sites or differences among species in dispersal ability to explore how mass effects, or source-sink dynamics, affect meatacommunity dynamics.

In his review of metacommunity concepts, Vellend (2010) portrayed community ecology as a "black box" that focuses more on pattern than process, reiterating Lawton's (1999) criticism that community ecology remains a "mess." To simplify the vast complexity of potential models and clarify similarities and differences among concepts and theories, he proposed that ecologists analyze community assembly as the outcome of four fundamental processes: selection, drift, speciation, and dispersal. Selection reflects the deterministic action of species sorting and filtering along environmental gradients and competitive interactions within habitats – the traditional domain of plant ecology and two of the four mechanisms in both Schmida and Wilson's (1985) and Leibold et al.'s (2004) schemes. Drift refers to the stochastic forces acting on population and community dynamics, driving some species to higher abundance and others to extinction. Speciation also affects community diversity but generally over periods of thousands of generations. Dispersal represents processes acting among sites that are fundamental to mass effects and all metacommunity approaches (Holyoak et al. 2005). Dispersal combines with local selective ecological processes to create a "vast" range of potential outcomes (Vellend 2010).

Given this rich body of work on metacommunity theory and models, we can ask how successful has this work been for predicting species gains and losses (turnover) and shifts in abundance? Such predictions would be particularly valuable to conservation biologists concerned with knowing where to focus their limited resources to conserve species most efficiently.

Higgins *et al*. (2006) and Azeria and Kolasa (2008) both suggested using nestedness in metacommunity matrices to predict future colonizations and extinctions. Azeria and Kolasa (2008) had some success, finding that extinctions of invertebrates in tropical rock pools decreased with predicted occupancy. In contrst, Azeria *et al*. (2006) found extinction probabilities for birds in the Dahlak archipelago to peak at intermediate occupancy probabilities. Donlan *et al*. (2005) found that community nestedness patterns failed to predict historical Holocene extinctions of mammals, leading them to conclude that tools developed from biogeography principles should be "evaluated critically" before being used in conservation planning.

Others appear even more disappointed at prospects for using metacommunity patterns to predict species turnover. Keith *et al*. (2011) recorded changes in a southern English woodland metacommunity across 86 ancient semi-natural woodlands over a 70-year interval. They found metacommunity structure to be stable despite declines in beta diversity and concluded that "metacommunity structure would not be a good landscape-scale indicator for conservation status." Others seeking to apply metacommunity models also express frustration in trying to predict patterns of species loss and colonization. Fleishman *et al*. (2002), in analyzing bird and butterfly occurrences, conclude that the factors influencing their distribution differ from place to place and among taxonomic groups, preventing us from using results from one group as following a general patterns that could apply to other groups. Fischer and Lindenmayer (2005) also found quite different patterns in studying the effects of fragmentation on the distributions of birds, arboreal marsupials and lizards in Australia. This led them to recommend autoecological studies of particular taxa over approaches based on metacommunity patterns. In studying beetle metacommunities in Tasmania, Driscoll (2008) also found that only certain subsets of the fauna followed any particular metacommunity model and that only about a third of the species showed evidence of deterministic metapopulation patterns. Driscoll and Lindenmayer (2009) went even further to assess predictions from six different theories against three classes of data on bird and reptile distributions over hundreds of sites in Australia. They found little consistent support for any of the theories as different species responded differently (and often temporarily) to differences in environmental conditions and geographic distance. Reflecting on these complex responses, they conclude that "metacommunity ideas cannot yet be used predictively in a management context." Finally, Lessard *et al*. (2012), after comparing local ecological processes among regions and highlighting the dangers of not considering differences in species pools when assessing the relative importance of species sorting and ecological filters, conclude that "there is no ‘proper’ scale with which to delineate the species pool, because species pools are shaped by multiple processes operating at different spatial and temporal scales, each of which can influence local patterns and processes." This represents a thorny issue and a serious criticism of previous efforts to analyze metacommunity dynamics.

The models we develop here accept the complexity and ambiguity inherent in trying to analyze metacommunity dynamics. Rather than trying to penetrate the community "black box" to dissect the several mechanisms at work, their scales of action, and their relative strengths and interactions, we instead capitalize on the rich information inherent in the metacommunity matrix itself to make predictions that we then rigorously test against empirical data. In that sense, our models resemble other classic approaches to analyzing ecological patterns in that they consciously ignore the complexities of species interactions and species responses to local site conditions. Simplifying assumptions, stochastic models, and dispersal are central to many theories of community organization that nevertheless have proved useful when tested against empirical data (Diamond 1975, Whittaker 1975). In introducing their theory of island biogeography, MacArthur and Wilson (1967) deliberately treated species as ecologically equivalent and assumed that islands differ only in area and isolation. Despite these simplifying assumptions, their theory continues to yield remarkably accurate predictions for species numbers and turnover across a huge number of archipelagos and fragmented terrestrial habitats (e.g., Newmark 1987). The simple models from island biogeography and Bell (1991) and Hubbell's (2001) neutral theory also serve as reliable points of departure for building more elaborate models that incorporate more deterministic mechanisms like species differences affecting how they sort along environmental gradients, interact with other species (competition, herbivory, disease, etc.), and disperse across complex landscapes.

Here, we develop models of metacommunity dynamics to predict changes in species occurrence and abundance within particular sites based only on the metacommunity matrix. These models present certain advantages including simplicity, the need for no initial data beyond the metacommuity matrix, and that they make clear testable predictions. They resemble other simple models like those reviewed above by intentionally ignoring many details known to affect ecological patterns and outcomes. These include differences in the ecological characteristics of species (beyond their incidence or abundance) and differences in site condition or location (beyond site richness or total plant abundance). Our models thus most obviously reflect the action of mass effects and dispersal in that they use each species' regional prevalence or abundance (row sums in the matrix) to make their predictions and in assuming that these affect species' local incidence or abundance. In addition, the models reflect some effects of deterministic factors like species sorting or competition/herbivory/disease in that they use information on how species richness (or total plant abundance) varies across sites (the column sums). Aside from these implicit (and assumed additive) effects, our models ignore mechanisms and ecological details including species characteristics, site effects, proximity and landscape effects, and all potential interactions among these. Our models differ from island biogeography in taking no account of island or patch area or distance. Like neutral models, our models predict no overall shifts in incidence or abundance. Unlike neutral models, our models predict directional changes in species incidence and abundance for individual sites rather than identical random walks for all species and sites.

To evaluate these models, we test their predictions against long-term changes in plant species incidence and abundance observed across 156 forested sites in southern and northern Wisconsin, USA in the 1950s and 2000s. These extensive datasets provide considerable power and a long interval for testing predictions from these models. We also formulate randomized null models to ensure fair tests of the models' predictions and to confirm that our results do not reflect any artifacts of model assumptions or the structure of these data. Finally, we use over-sampled data from the northern sites to assess how limiting sampling to fewer quadrats or the most abundant species can affect the accuracy and power of the models' conclusions. Together, these efforts demonstrate both the power and limitations of this approach. Finally, we discuss how such models might be used both by ecologists seeking to understand the particular forces affecting community dynamics and by conservation biologists eager to predict site-specific population dynamics.

## Methods

### Metacommunity incidence and abundance models

The two models we present are identical in structure and assumptions but make complementary predictions regarding species dynamics. The incidence model outlined in Fig. 1 uses only data on species' presence and absence to predict expected changes in species incidence via local colonizations and extinctions. In contrast, the abundance model predicts changes in species abundance but only for those species that persist at sites. Both models use the products of row and column totals in the metacommunity to make predictions. In using products of the marginal sums in these models, we clearly assume that:

1. The likelihood that a species will occupy a site can be estimated from the product of its overall prevalence across the metacommunity and the richness of species at that site.
2. Changes in the abundance of a species persisting at a site can be estimated from the product of its overall abundance across the metacommunity and the total abundance of all species occurring at that site.
3. These effects of species prevalence (or abundance) and site richness (or total plant abundance) are additive and thus do not interact.
4. Any mechanisms acting to change patterns of species incidence (or abundance) across the metacommunity in these models must act through these row and column totals.

**Figure 1.**
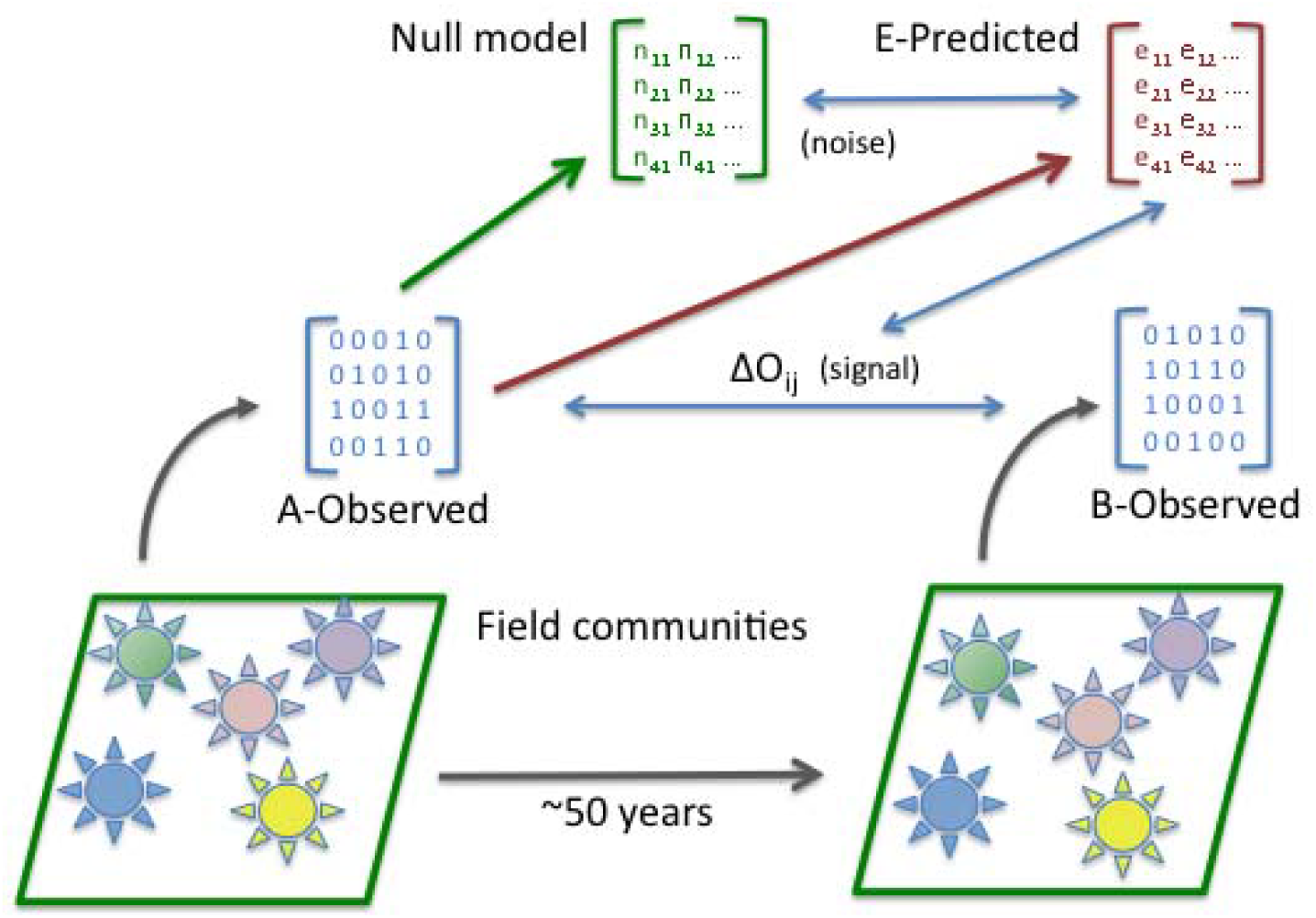
Modeling process. The diagram shows steps in the analysis. Data on species incidence (shown here) or abundance obtained from the metacommunity at an initial time A are used to generate a metacommunity matrix (A-Observed). The row and column totals from this matrix, in turn, are used to generate a matrix of predicted values (matrix E-Predicted with values Eij) for either expected occupancy (the probability that species i will occur at site j) or their expected abundances. Values from matrix A-Observed are also used to generate 100 matrices with randomly shuffled values subject to certain constraints (see text). These represent expected values if changes in the metacommunity occur randomly (the Null model) rather than according to the incidence or abundance predictive models presented.

Thus, we acknowledge that the marginal sums are influenced by several factors, possibly including local site favorability, environmental and biotic filtering, landscape connectivity, dispersal limitation, and ecological drift. Although none of these factors appear explicitly in the models, they likely incorporate both mass effects and some combination of these other factors. We see the implicit structure of our models as a strength, however, in that they require no explicit information on any of these processes and make no assumptions about their relative importance or how they combine and interact. Rather, the models implicitly integrate effects of these mechanisms via the row and column totals to generate their predictions.

A metacommunity can be represented by a matrix containing data on either species incidence or estimates of species abundance (Fig. 1). Species are generally arranged as rows and sites as columns. Cells within an incidence matrix, **O**, reflect the presence or absence of species at sites with O_ij_ = 1 if species *i* was observed at site *j* and 0 otherwise (Fig. 2a). Cells within an abundance matrix, **F**, reflect species abundances with F_ij_ equal to the observed or estimated abundance of species *i* at site *j* (Fig. 3a). We develop and test models for both types of matrix. Our incidence model predicts the likelihood that any given species will occur at any given site from the product of its row and column totals. In particular, the expected probability that species *i* will occur at site *j* is:

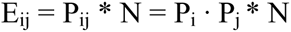

where P_i_ is the row sum for species i divided by the matrix sum, P_j_ is the column sum for site j divided by the matrix sum, and N is the matrix sum (an example appears in Supplementary Table S-1a). Note that this model weights the likelihood that any given species will occur at a given site identically by both its overall incidence across all sites and by the number of species that occur at that site (matching how Chi-squared tests calculate expected values in contingency tables). Expected values for the *abundance* model (EF_ij_) are calculated in the same way except that the weightings reflect products of each species' total abundance across all sites (row sum F_i•_) and the total abundance of all species within each site (column sum F_•j_ – Fig. 3a) so that EF_ij A_ = F_i•_ · F_•j_/F_tot_, where F_tot_ is the total abundance summed across species and sites (Fig. 3C).

**Figure 2.**
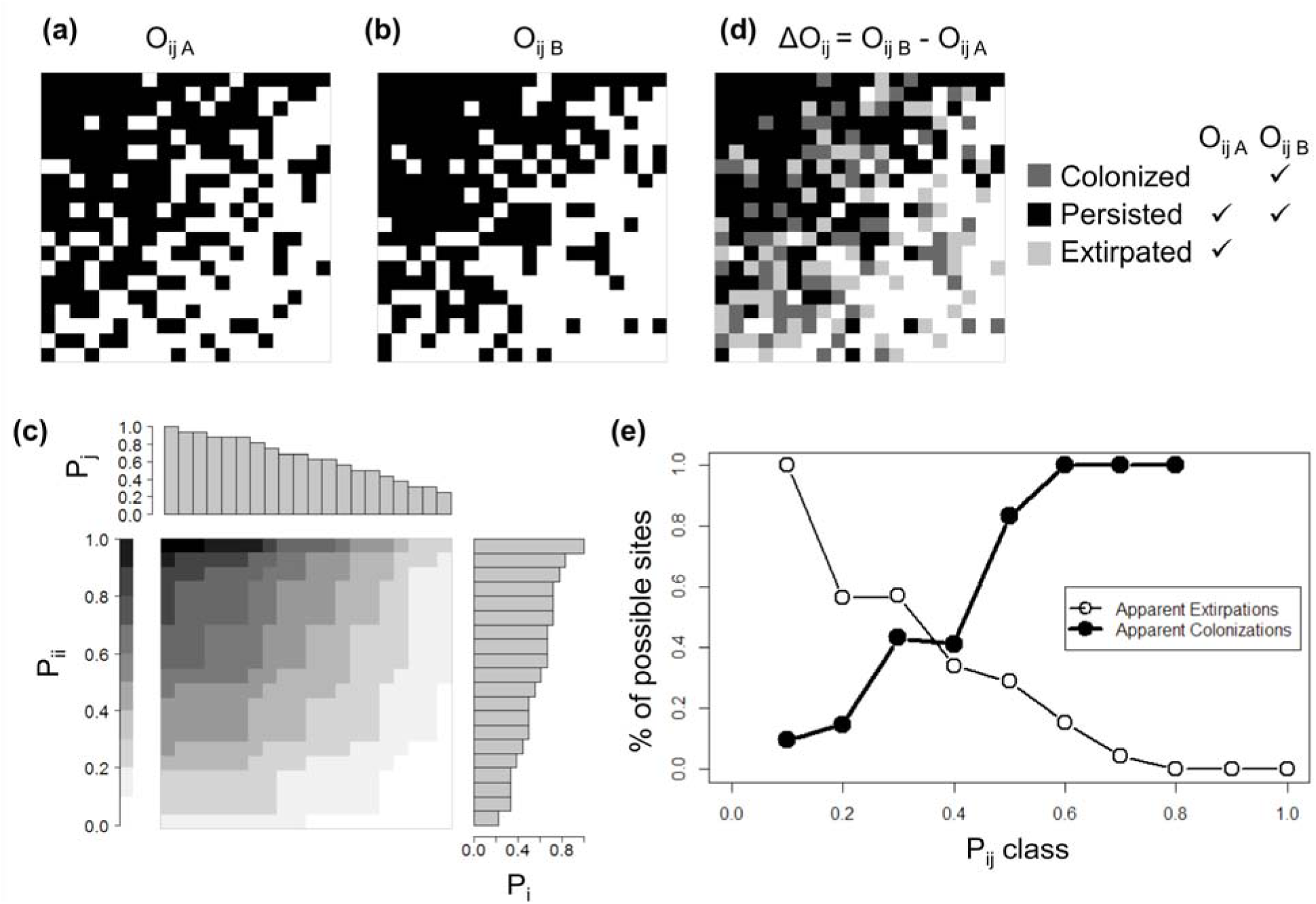
The incidence model. (a) An observed species incidence (presence/absence) matrix, O_ij A_, in initial year A based on fictional data representing a small number of species that are relatively common but differ in abundance as they might occur across a small set of sites that differ in diversity. Cells are black if species *i* was observed at site *j*, and white otherwise. We sort the matrix by row totals (O_i•_ or the number of sites where species *i* was observed = total species incidence) and by column totals (O_•j_, or the number of species observed at site j = site richness) but this is not a necessary part of the model. (b) The observed presence/absence matrix for Year B, O_ij B_, again sorted by row and column as in (a). (c) Expected probabilities of occupancy E_ij_ = P_i_ · P_j_, ^*^ N where *P* is the proportion of sites occupied by species *i* (bars to the right), P_j_ is the proportion of species in the dataset found at site j (bars at the top), and N is the overall matrix sum. Cells are shaded by prediction bands, rounded to the nearest tenth. (d) Matrix showing local extinctions, colonizations, and persistence between periods for the two incidence matrices.(e) Proportion of available sites at each probability of occupancy class that were either colonized between periods (filled circles) or experienced a local local extinction (open circles).

**Figure 3.**
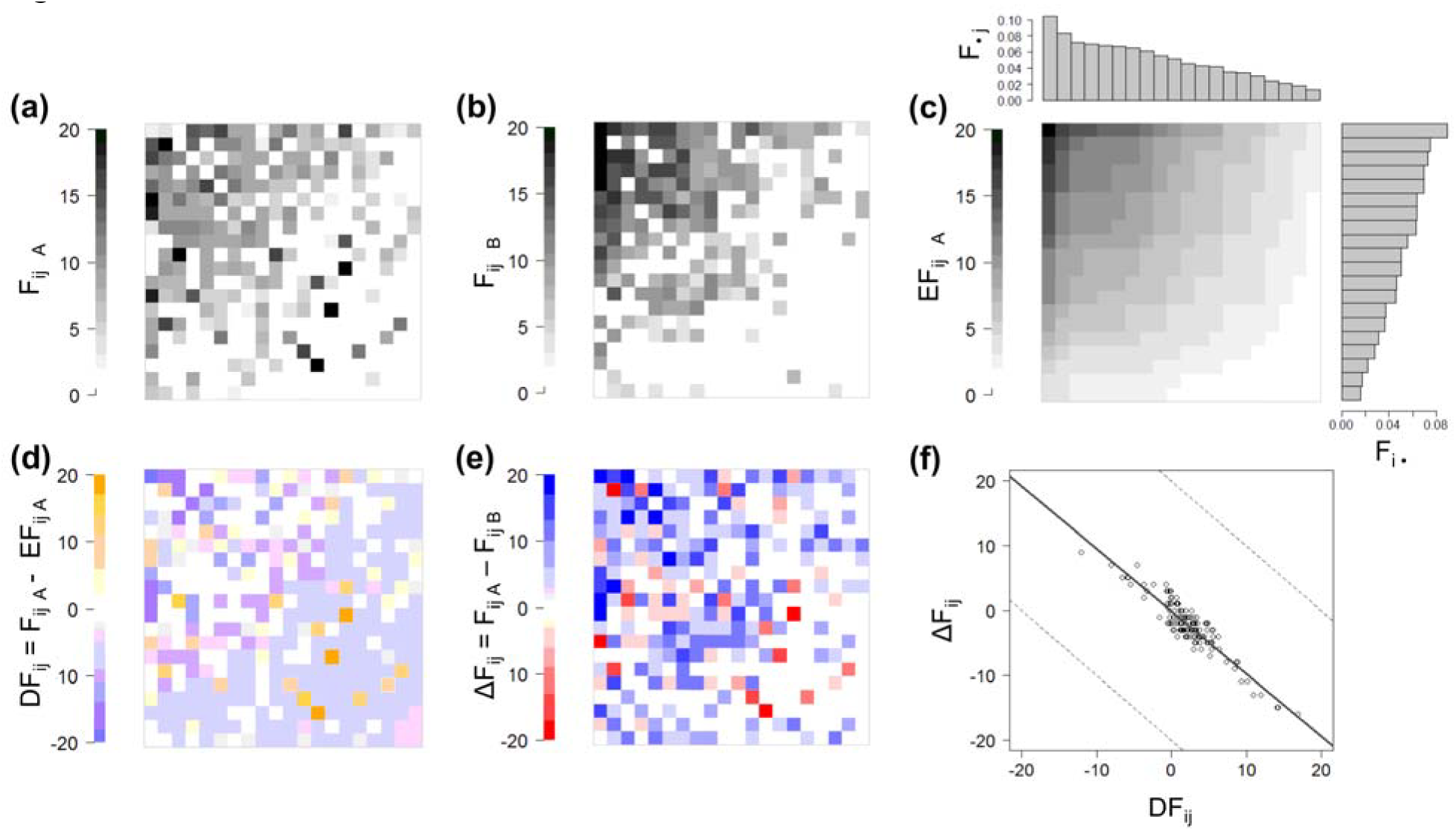
The abundance model. (a) F_ij A,_ observed frequency matrix in initial year A (fictional data). (b) F_ij B_, observed frequency matrix for Year B. (c) Expected frequency for species i at site j for Year A, EF_ij_ = F_i•_ ^*^ F_•j_ ^*^ F_tot_, where F_•j_ and F_i•_. are as above, and *F*_*tot*_ is the sum of all quadrat occurrences across species and sites. (d) Change in frequency over the two time periods, ΔF_ij_ = F_ij B_ – F_ij A_. (e) Departures of observed from expected frequencies in each matrix cell, DF_ij_ = F_ij_ – EF_ij_. (f) Comparing matrix cell values of ΔF_ij_ to those of DF_ij_. The solid line shows the linear best fit while dotted lines reflect limits on the values possible due to sampling 20-quadrats. Matrices are sorted by row and column totals to aid visual interpretation.

Field surveys at an initial time A generate the data used to compute the original metacomunity matrix (**O** or **F**) that is then used to predict these expected values (matrix **E** – Fig. 1). These expected values are then compared to a second observed metacommunity matrix derived from re-surveys of the same sites at a later time B. A difference matrix is then computed between the two successive observed incidence or abundance matrices with elements:

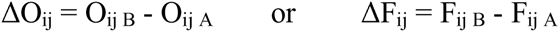

(Fig. 2d). Values in the incidence difference matrix can be 1 (reflecting colonization of a previously unoccupied site), -1 (local extinction), 0 (persistence), or NA (cells where a species did not occur at either time A or B). The analogous abundance difference matrix records changes in observed abundance. These are most meaningful for species present at both times A and B at a site.

Note that no set time period is explicitly assumed in these models. To make useful predictions, the time period needs to be long enough to allow biological turnover, but not so long that major disturbances occur, restarting the successional clock, or that great changes in community composition or abundance occur. Because predictions of the models depend only on conditions at a single previous time, these are Markov models (Feller 1968).

Note also that these models predict no overall changes in incidence or abundance – colonizations and increases in abundance are balanced by local extinctions and declines (see row and matrix sums in Table S-1). Initially rare species remain rare, common species remain equally common. Sites also retain the same overall richness or total abundance that they had originally (evident in the stable column totals in Table S-1). Most real metacommunities violate this assumption including the ones we use to test these models. Nevertheless, this assumption is parsimonious in not assuming or predicting any systematic changes in abundance or species richness. The models could easily be adapted to incorporate systematic changes in overall incidence or abundance for species, sites, or overall by multiplying all predicted values by the observed shifts in row, column, or matrix totals, respectively. These models differ from neutral models in not assuming local random increases or decreases in abundance or incidence that might change overall abundance (see Neutral model matrix, Table S-1).

### Testing predictions of the models

These models make specific predictions. For the incidence model, species that do not occur at sites at time A where they are expected to (E_ij_, > O_ij A_), we expect colonizations may occur. For species that occur at sites where they are not expected to at time A (E_ij_ < 1), we expect appreciable rates of local extinction. Likewise for the abundance model, species already present at a site will likely increase in abundance there when their abundance at time A is less than expected (F_ij A_ < E_ij_) and vice versa. Specifically, we calculate differences between the actual observed initial frequencies and those expected on the basis of row and column sums :

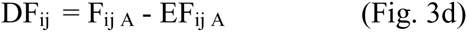

Positive values of DF_ij_ reflect cases where species i at site j had a higher abundance than expected at time A under the model while negative values reflect the opposite.

We evaluate these predictions for the incidence model by binning values of E_ij_ into ten intervals and examining how the proportion of local extinctions and colonizations varies as these values of E_ij_ increase (Fig. 2e). We quantify the strength of these trends by fitting a slope to these values via linear regression. Abundance model predictions mirror those of the incidence model but instead predict how species that persist at a site are likely to shift in abundance. As noted, the expected abundance at time A, EF_ij A_, reflects the product of the abundance of species *i* across all sites at time A (row sum F_i•_) and the abundance of all species at site *j* (column sum F_•j_ – Fig. 3a). Specifically, EF_ij A_ = F_i•_ · F_•j_/F_tot_, where F_tot_ is the total abundance summed across species and sites (Fig. 3c). We then generate predictions from this model by calculating differences between the actual observed initial abundances and those expected on the basis of row and column sums: DF_ij_ = F_ij A_ - EF_ij A_(Fig. 3d). Positive values of DF_ij_ reflect cases where species i at site j had a higher abundance than expected at time A while negative values reflect the opposite. To test the model, we compute the observed difference matrix reflecting how species that persisted at each site shifted in abundance over the interval: ΔF_ij_ = F_ij B_ - F_ij A_ (Fig. 3e). We expect ΔF_ij_ to decline with increases in DF_ij_ allowing us to assess predictions of the model using linear regression (Fig. 3f).

### Null model randomization tests

To provide reliable statistical tests of these predictions, we created null models to estimate the random shifts in incidence or abundance that might be expected in metacommunities experiencing no systematic local extinctions, colonizations, or shifts in abundance (Fig. 1). Null models for static species x site matrices exist (Ulrich 2007, Ulrich and Gotelli 2010), but null models for community change do not. To create a null model for species incidence, we separately randomized species occurrences across rows in proportion to each species' overall incidence in the metacommunity (P_i_). In particular, for each species, we first shuffled observed values (the 0's and 1's for that species) among the cells that lacked that species in the original row (i.e., where O_ij A_ = 0). We restricted changes in these cells to be either a colonization event (0 ➔ 1) or never present (NA). We then performed a similar randomization over the initially occupied cells in that row (O_ij A_ = 1), restricting these changes to reflect either a local extinction (1 ➔ 0) or persistence (1 ➔ 1). We imposed no row or column total restrictions on these randomizations or on the resulting presence/absence matrices. Imposing such restriction would bias the resulting pattern making it less than fully random.

To test for significant departures of the observed incidence data from the randomized null model, we compare slopes of lines fit to the observed data (signal) to those fit to the null models (noise) for both local extinctions and colonizations (Figs. 1 and 2e). The slopes of these best-fit lines provide a statistic to evaluate trends in local extinction and colonization across successive P_ij_ classes. If the slopes fitted to the observed data are steeper than 95% of the slopes derived from the null model, we interpret the observed changes as systematic.

We test predictions from the abundance model using a similar null model. Specifically, we compared the relationship between the observed changes in abundance, ΔF_ij_, to the changes expected from random processes, DF_ij_ (Fig. 3f). To generate random values for these null model matrices (ΔF_ij-sim_), we shuffle all the observed values for a species among the subset of cells that that species occupied in either sample period leaving other cells at zero. This procedure assumes that species only occur at sites where they were observed and that changes in a species' abundance are random and independent of the abundance of that species in the landscape and the total abundance of species found at any given site. This random shuffling could assign a large decrease in abundance to cells originally at low frequency, resulting in a negative frequency. Similarly, a large increase in abundance might be assigned to cells already at high frequency. To prevent this, we constrained abundance between a floor of 0 and a ceiling set by its maximum possible value (a frequency of 20 in the empirical tests described below). We then used the correlation coefficient and slope of the best-fit line between the observed and predicted changes (ΔF_ij_ and DF_ij_) as statistics to compare with analogous values generated via the null models.

### Field data

We use empirical data on the occurrences and abundances of understory plants across 156 forested sites in southern and northern Wisconsin (Fig. 4) to test predictions from these models. These two regions differ conspicuously in climate, soils, and landscape context and conditions causing their forests to differ considerably in type and composition. They occupy distinct floristic provinces separated by a well-recognized “tension zone” (Curtis 1959). The northern and eastern portions of the state contain hardwood and coniferous forests while the southern and western portions supported prairies and hardwoods (Curtis 1959, Curtis and McIntosh 1951). Forests remain dominant in northern Wisconsin whereas agriculture and periurban areas now dominate southern Wisconsin. There, the landscape has 12-15x higher densities of roads, people, and housing than the northern forests (Radeloff *et al*. 2005, Riiters and al. 2002). These differences emerge conspicuously in multivariate analyses that show a perfect primary split in a cluster analysis based on species composition (incidence) and cleanly separated clouds in ordination space based on abundance (Fig. S-1). Because they are geographically removed from each other, the potential for seeds in one region to disperse to the other is limited. Given this isolation and these conspicuous differences in forest type, landscape context, and floristic composition, we treat the northern and southern sites as separate metacommunities.

**Figure 4.**
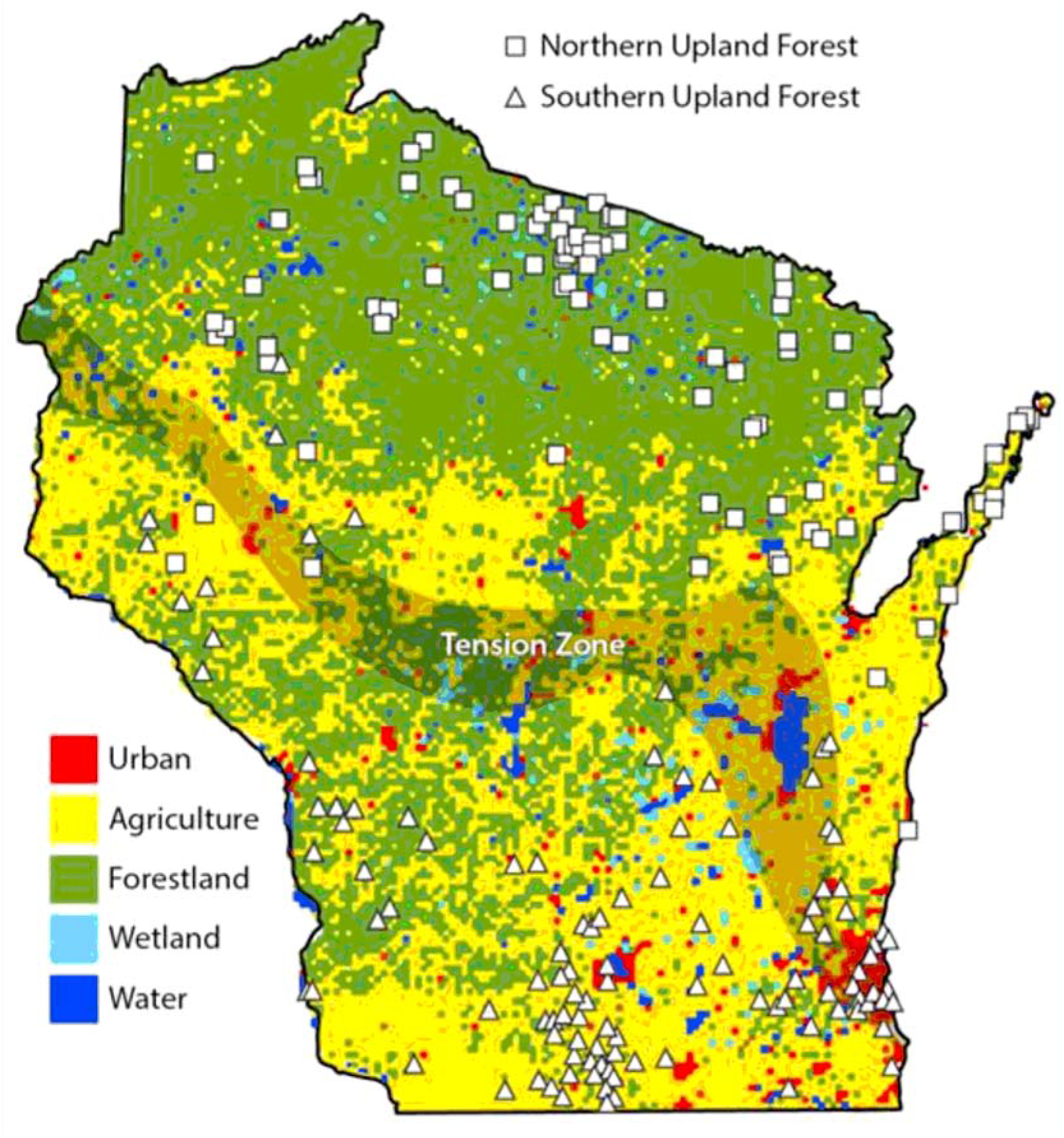
Forest site locations. Map showing the 62 sites in northern Wisconsin (squares) and 94 sites in southern Wisconsin (circles). The map also shows land cover across the state demonstrating the difference between the primarily forested landscape matrix in northern Wisconsin and the far more fragmented forests in southern Wisconsin (cover map from http://dnr.wi.gov/maps/gis/datalandcover.html).

Ideal metacommunties for testing our model would consist of a relatively large species pool distributed among many sites occurring across a contiguous and relatively homogeneous region, all sampled thoroughly using standard methods. Our system approximates this case. To allow adequate time for colonization and extinction events and changes in abundance to occur, the metacommunity should be resurveyed after a span of time sufficient to allow some turnover (several generations). Most of the species in these metacommunities are herbaceous perennials with lifespans of 5-25 years, allowing turnover over the 50-year period we use.

For the initial baseline (time A), we use high-quality legacy data collected between 1947 and 1956 (henceforth "1950s") by the Wisconsin Plant Ecology Laboratory under the leadership of J.T. Curtis (see http://www.botany.wisc.edu/PEL/). They developed quantitative methods to survey plant communities and applied these to hundreds of forested sites (Fig. 4) in both southern and northern Wisconsin. Curtis and colleagues sampled stands that were relatively undisturbed, uniform in topography, and >6ha in area. They characterized the overstory at these sites using plotless methods and sampled the understory by recording all vascular species present in each of 20 replicate 1 m^2^ quadrats spaced every 20-30m around a large square covering about 1 ha. They estimated plant abundance as the frequency of each species among these 20 quadrats. We then resurveyed many of these sites in the 2000s using similar but more intensive methods, again estimating abundance as the frequency at which species occurred among sampled quadrats (80 to 120 in these resurveys). We only resurveyed sites lacking recent disturbance and changes in land-use, placing new surveys as close as possible to the originals (within 50m). Wiegmann and others resurveyed 62 sites in northern Wisconsin in 2000 and 2001 using six 20x1m strip transects, resulting in 120 sampled quadrats (Rooney et al. 2004, Wiegmann and Waller 2006). Rogers et al. (2008, 2009) resurveyed the 94 southern Wisconsin sites in 2004 and 2005, sampling 80 spaced quadrats per site. To match sample sizes between periods, we subsampled the 2000s data using results from every sixth quadrat in the northern forests, creating six replicate samples each containing 20 spaced quadrats with similar spacing as the original surveys. In the southern forests, we used every second quadrat from the sampled square judged closest to the original survey site, again providing samples of 20 quadrats of similar spacing and extent as the original survey. We synchronized species lists between periods to reflect changes in nomenclature and possible misidentification, lumping taxa in a few cases. Species nomenclature follows Gleason and Cronquist (1991).

### Applying the model

We applied both the incidence and frequency models to both the southern and northern metacommunities using data from the 1950s surveys to generate predictions for subsequent shifts in incidence and abundance (Fig. 1). We first trimmed the full data set of rare species. Rare species present a potential source of bias in that they are easy to miss in limited samples leading to overestimates of both colonizations and extinctions. To reduce this bias and ensure more accurate abundance estimates, we only include species that are above the median (top 50%) in regional (metacommunity) abundance. This resulting metacommunity matrices contain 64 and 100 species distributed among 62 and 94 sites in northern and southern Wisconsin, respectively. We assessed the effect of using this threshold on our results via simulations that repeated our analyses over higher minimum abundance thresholds in the northern dataset requiring a total frequency in both periods of at least 0.005, 0.01, or 0.05 (Table 1). To confirm that our tests reveal true changes rather than pseudo-colonizations and extinctions from limited samples, we also used the six sets of replicates from the 2000s resurvey in N Wisconsin to simulate the sampling variance generated by resurveying the same plant communities using different samples. We then used each sample to generate a prediction of the "change" expected at the other five.

**Table 1.** Number of species retained for analysis after removing rare species at each of three cutoff levels.

|  | Northern |  | Southern |  |
| --- | --- | --- | --- | --- |
|  | (62 sites) |  | (94 sites) |  |
| Cutoff | 1950s | 2000s | 1950s | 2000s |
| 0.05% | 77 | 89 |  |  |
| 1% | 58 | 65 | 98 | 100 |
| 5% | 26 | 26 |  |  |

We then compared the fits from these 30 pairs ("noise") to the six independent fits of our model predictions to the 2000s data ("signal") for both the incidence and frequency models using these three frequency cutoffs. The outcomes of these simulations assure us that the results we present are robust and not a sampling artifact.

## Results

Species incidence and abundances in both forest metacommunities remained broadly similar between the 1950s and 2000s (Fig. 5). The mean incidence of species declined between the 1950s and 2000s in both regions even as the total frequency of plant species (our index of abundance) increased (Table 2). Overall species richness increased in southern sites but decreased in the North while mean frequency did the reverse. In northern Wisconsin, overall patterns of relative species incidence and frequency remained stable between the two periods (autocorrelations of *ρ* = 0.722 and *ρ* = 0.782, respectively, both *P*<0.001). Species incidence and frequency were less auto-correlated in the more dynamic southern forests (*ρ* = 0.516 and *ρ* = 0.489, respectively, *P*<0.01). Despite this stability, these communities declined in diversity with local plant richness per site, based on 20 quadrats, down ~15% in northern and ~25% in southern forests. They also shifted considerably in composition (Rooney et al. 2004; Rogers et al. 2009, Rogers et al. 2008).

**Figure 5.**
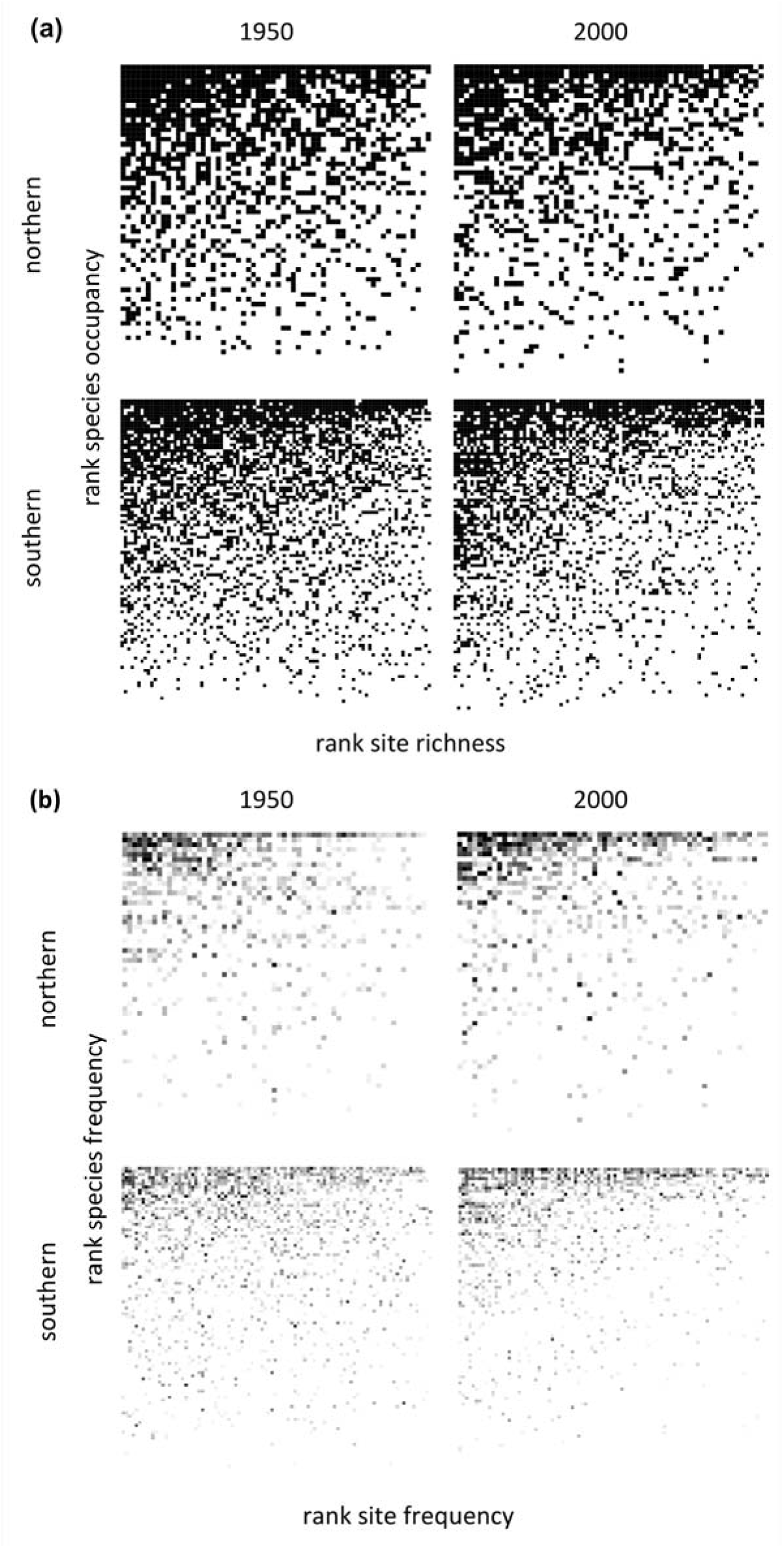
Species x site metacommunity matrices. (a) Sorted presence-absence matrices (Oij) for sites in northern (showing 1% cutoff) and southern WI in both the 1950s and 2000s. Matrices are sorted by row totals (the number of sites each species occupies) and column totals (site richness) within each metacommunity. (b) Analogous frequency matrices (OFij; sorted by row and column totals). Cells colored darker gray indicate more quadrats occupied at a site. Southern matrices appear finer-grained as they reflect both more site and more species.

**Table 2.**
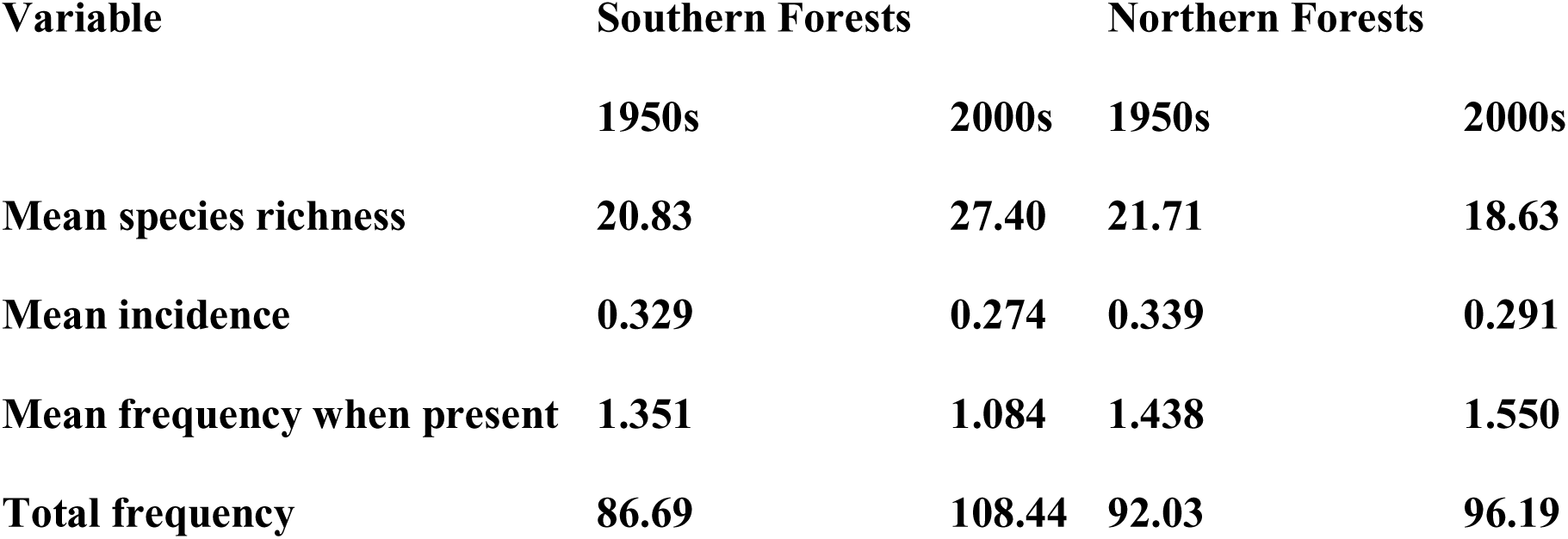
Summary statistics for the Wisconsin forest plant community samples. Values here reflect the means across sites calculated from the metacommunities used here. These values pertain only to the more abundant species included here.

The incidence and the abundance models both served to predict changes in these communities across the 50-year interval (Fig. 6). Predictions from the incidence model paralleled observed changes in site occupancy with occupancy and colonizations increasing and local extinctions declining as the model predicted they would (Fig. 6a, b). These trends are all highly significant in Cochran-Armitage tests (all P<0.001). In fact, in the richer southern forests, the best-fit slope to the observed vs. predicted trend in colonizations was a remarkable 0.94, far higher than any of the 100 slopes generated by the null model (range: -0.23 to 0.58, with a mean of -0.034; Fig. 6b). Observed local extinctions also declined sharply as predicted occupancy increased (slope = -0.66) whereas the slopes generated by the null model range from 0.28 and +0.23 (mean: -0.0083; Fig. 6b).

**Figure 6.**
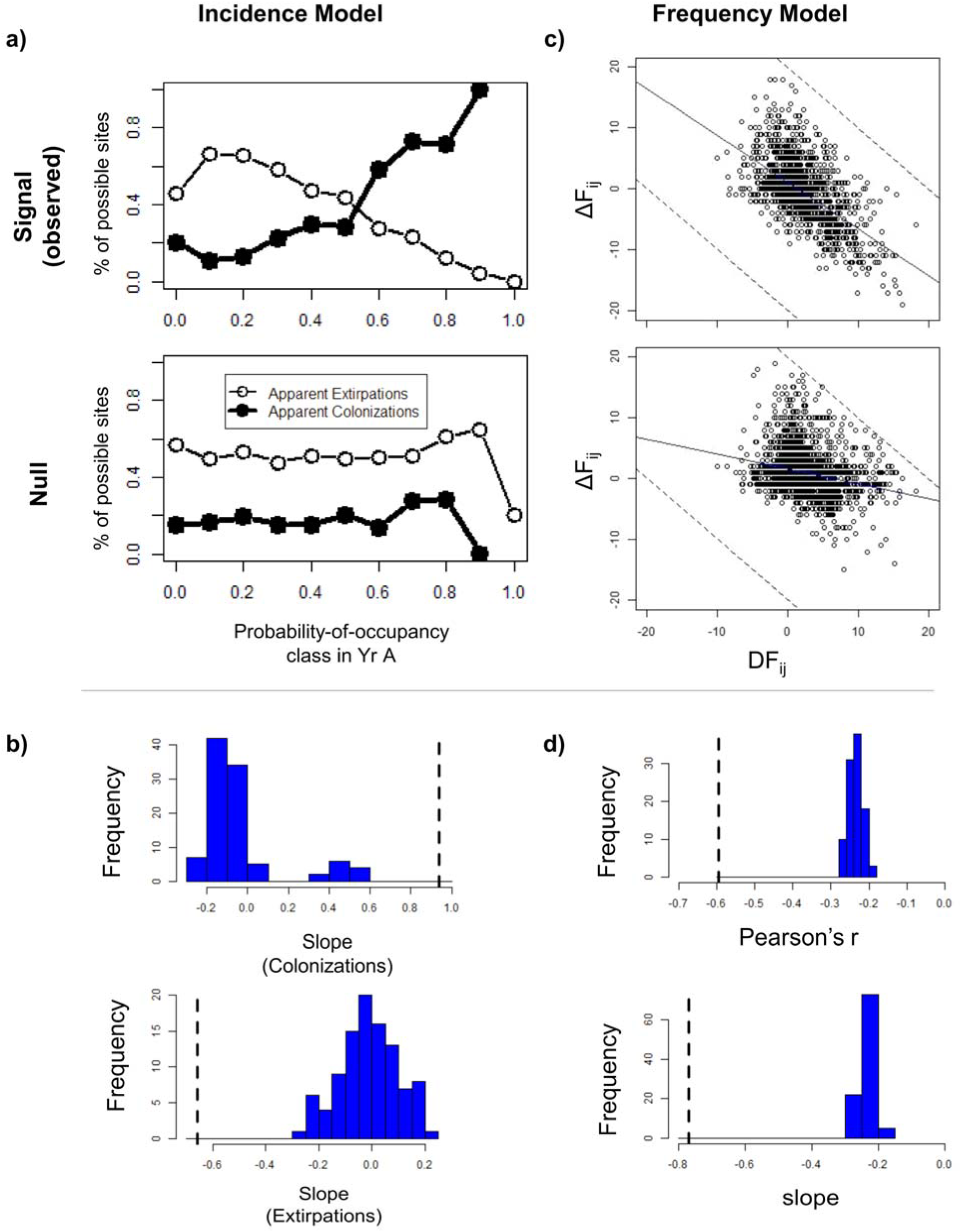
Tests of the incidence and frequency models. Results shown reflect tests on the larger data set from southern Wisconsin. (a) The observed (“signal”) and one example of a “null” simulation of the local extinction trends (empty circles) and colonization trends (filled circles) across P_ij_ classes. (b) Histograms of values for the slope of the best-fit line to the colonization and local extinction trends for the 100 simulations of “null” data. The value for the observed “signal” is shown with a heavy dotted line. (c) Scatter plot of AF_ij_ vs. *DF*_ij_ for the observed (“signal”) and one of the simulated null model data sets. (d) Histograms of values for the Pearson’s correlation coefficient *r* and slopes of the best-fit lines between ΔF_ij_ and *DF*_ij_ for the 100 simulations of the null data sets. The value for the observed “signal” is shown with a heavy dotted line.

The occupancy model also served to predict metacommunity colonizations and extinctions in the northern forests (Fig. S-2 a). Actual colonizations increased sharply in cells with higher predicted probabilities of occupancy with all slopes exceeding those generated by any of the null models using species with frequencies above 0.5% or 1% (Fig. S-2 a and b). The model's predictions weakened, however, when rarer species (with a frequency <5%) were excluded (Fig. S-2c). Actual local extinctions also declined strongly as predicted occupancy increased. These mean slopes decreased only slightly (from -0.63 to -0.61) in moving from the 0.5% to the 1% frequency threshold and remained highly significantly different from the mean slopes for the null models (0.012 & 0.003). Again, the pattern weakened when all species with frequencies below 5% were excluded.

The abundance models had even more predictive power than the incidence models with observed changes in species frequency over the past 50 years tracking predictions of the models (Figs. 6c and S-3a). Species locally more abundant in 1950 than expected in the model tended to decline in abundance and vice versa (signal biplots). These slopes (ΔF_ij_ vs. *DF*_ij_) were again far steeper than those of the null models (Fig. 6d and S-3a). In the southern forests, the correlation was -0.60 and the slope -0.77, again far exceeding those generated by the null models (r: -0.28 to -0.19, slopes: -0.28 to -0.18; differences all P<0.01, Fig. 6d). Predictions for abundance in the northern forests showed similarly dramatic differences from the null models and these differences persist even when all species with a frequency below 5% are excluded (Fig.S-3).

In applying these models, we pruned the rarest species in order to reduce the influence of random fluctuations. However, pruning species also reduces sample size, potentially limiting our ability to test these models. Because most species are rare, increasing the minimum frequency necessary to be included in the analysis from 0.05% to 1% reduced the number of species by 25-27%. Increasing the threshold to 5% (including only common species) dropped sample size far more – from 77 (89) species in the N (S) Wisconsin metacommunity in the 1950s (2000s) to just 26 in both. Despite including a third or fewer of the species, these drops in the threshold hardly affected the signal (or noise) in the abundance model. The observed change in frequency continued to closely track model predictions (r = -0.46, -0.46, and -0.45) across all three abundance thresholds, always exceeding correlations in null models (means: -0.20, -0.21, -0.22, Fig. 7a vs. 7b vs. 7c). Likewise, ΔF_ij_ tracked the *DF*_ij_ values closely (signal means: -0.67, -0.67, and -0.66 for the three thresholds), and again far exceeded values observed in the null models (means: -0.22, -0.23, -0.25) or in the replicated spatial noise data (means: -0.084, -0.082, -0.082). We conclude that the abundance model (based here on species frequencies) is quite robust to partial sampling that only includes more abundant species. It is also robust to spatial variation in sampling, providing high signal to noise in all cases. Thus, the models' success in predicting the changes actually observed among these communities is real and neither an artifact of sampling common species nor of pseudo-colonizations and extinctions generated via incomplete sampling.

In contrast, raising the species abundance threshold reduced power in the incidence model – as might be expected given the fewer species included. Null model predictions remained stable but the observed signal declined (Fig. S-2a vs. b and c). Pruning rare species greatly reduces sample sizes for estimating colonizations and local extinctions. The highest abundance threshold (a frequency of >5%) excludes over two thirds of the species from the analysis. In this case, the local extinction signal slope (-0.31) remains somewhat greater than the null model slopes (mean -0.015) but colonization slopes (mean: 0.075) barely exceed these (means: 0.025 and 0.03; Fig 6c). We conclude that to use the incidence model, we need thorough sampling (enough to detect species at an overall frequency of 1% or less) to ensure model power.

We also used replicate samples from the northern forests to assess how sampling variation affects our estimates of local shifts in incidence and abundance. The observed local extinction slopes of -0.63 and -0.61 are far steeper those generated by the noise models (means: -0.29 and -0.28, Figs. S-2 & S-3, "noise" rows). Thus, the success of the incidence model does not reflect sampling artifacts. For colonizations, the patterns are similar except that slopes for the “noise” tests vary more, partially overlapping the “signal” slopes (Fig. S-2a, b colonization panels). The difference between the observed and noise colonization rates is greatest in the highest occupancy class, as expected given that the more abundant species provide more power to distinguish signal from noise.

## Discussion

The models of metacommunity change that we introduce and test here demonstrate the power that the information contained within the metacommunity matrix itself has for predicting changes in species incidence and abundance. The accuracy and significance of these predictions for the many sites and species in these two, largely distinct, metacommunities surprised us.These models contained no explicit information on species' functional or behavioral traits, nor any data on site conditions, proximity, or landscape context. Nevertheless, the models served to predict changes in plant species incidence and abundance over the succeeding 50 years in the forests of southern and northern Wisconsin. The abundance models accounted for 35% of the variation in observed shifts in frequency in the forests of southern Wisconsin (vs. <6% for the null model) and 23% (vs. 4%) in the northern forests. All four models generated predictions with high statistical significance showing no overlap with the predictions generated by the matched null models (Figs. 5 and 6). Given the many site, landscape, and species characteristics known to affect species and community dyanmics, it is remarkable to find so much predictive power in such a simple model.

Predictions from these models independently fit two quite different metacommunities that differ greatly in forest type, soils, climate, and landscape context. The abundance models had more predictive power than the incidence models. This should be expected given that abundance data contain more information than species presence / absence. Predictions for both the incidence and abundance models also proved to be more accurate in the far more fragmented oak-hickory forests of southern Wisconsin than in the continuous mixed deciduous / coniferous forests in the North. This was particularly true for predicted colonizations. This could reflect the higher number of species in the South, the greater number of sites, and/or the greater changes in landscape conditions that have occurred there, increasing species losses and turnover (Rogers et al. 2009, Rogers et al. 2008, Rooney et al. 2004). The resulting species losses reflect an extinction debt related to habitat area and isolation (Rogers et al. 2009). This may explain why the more continuous forests in northern Wisconsin show fewer shifts in the relative rankings of the row and column totals. It is also remarkable that the models succeeded well given that they assume no net change in local species richness while each site actually lost, on average, 15% or 25% of its species in the northern and southern forests, respectively (Rooney et al. 2004; Rogers et al. 2008).

The success of our models may reflect the quality and quantity of data available for these forests. The surveys and resurveys incorporated quantitative data for 64 and 100 species across 62 and 94 sites in northern and southern Wisconsin, respectively, over a 50-year period. Such extensive data provide reliable row and column totals and a biologically meaningful interval long enough to allow appreciable turnover. Communities sampled at fewer sites, with fewer species, or with fewer quadrats could reduce the accuracy of model predictions. Likewise, more closely timed surveys would show fewer and smaller changes. While we urge others to test these models in other systems, long-term resurvey data remain scarce (Waller and Rooney 2004, 2008).

Oversampling during the 2000s resurveys of the northern sites allowed us to assess re-sampling noise associated with the pseudo-colonizations and extinctions inevitable in resurvey data. These had minor effects relative to the strong signals from our models and the actual changes observed. The replicate sampling further allowed us to assess effects of including more or fewer rare species. These thresholds had little qualitative or quantitative effect on results from the abundance models, perhaps reflecting the fact that such models focus on species present at both sampling periods which tend to be common. In contrast, performance of the incidence models – particularly for colonizations – declined when we only used the 26 most abundant species. Given that colonizations and extinctions occur mostly among rarer species, this is not surprising. Additionally, focusing on more abundant species produces a fuller overall matrix, creating fewer opportunities for colonization than extinction. This may account for the fact that we witnessed stronger slopes for extinctions than for colonizations. Researchers eager to apply the incidence model should be cautious when applying it to metacommunities with few species.

The accurate predictions of these models in two distinct metacommunities suggest that these models may prove useful in additional regions. Alternatively, their success here might reflect a fortuitous selection of relatively undisturbed sites and a dynamic set of sites resurveyed over an ideal interval. Before concluding that they are general and useful, these models should be applied and tested in other contexts. Following several such tests, we could compare studies to identify particular conditions under which these models perform better or worse. For example, communities with more species, more sites, or longer intervals between surveys might provide better fits to these models. Our results comparing northern to southern sites and trimming species in the simulations suggest that predictions based on smaller metacommunities may be less powerful than those from larger ones, at least for the incidence model. Likewise, we predict that incidence models will be more sensitive to sample size than the abundance models.

These incidence and abundance models lack explicit information about species characteristics or site or landscape conditions, yet species gains and losses are hardly random with respect to the species or sites involved. Verheyen et al. (2004) found that species with low seed production and short-distance seed dispersal had lower rates of colonization and extinction in the forests of central England and were more likely to show effects of patch age and connectivity than other species. Such findings motivate the more complex models that ecologists often apply to metacommunities. These include information on species, sites, and/or (meta)community structure. Assembly rule models explicitly incorporate differences among species in dispersal, competitive ability, or other traits (e.g., Cornwell et al. 2006, Duckworth et al. 2000, Feeley 2003, Grover 1994, Messier et al. 2010). Successional and microsite models focus instead on site conditions and disturbance to predict community composition (e.g., Matthews 2004, Wethered and Lawes 2005). Community nestedness has also been used to predict changes in community composition (e.g., Azeria et al. 2006, Báldi 2003, Baselga 2010, Cook and Quinn 1995, Cutler 1991, Fischer and Lindenmayer 2005, Lomolino 1996, Maron et al. 2004, Patterson and Atmar 1986), as have patterns of species co-occurrence (e.g., Sfenthourakis et al. 2006). All these models are more elaborate than the models we present here in that they rely on specific data or assumptions regarding how species interact with abiotic or biotic conditions.

Our models most obviously reflect the action of mass effects in using each species' regional abundance (the row sums) to make predictions. That the models work well suggests that mass effects are an important part of metacommunity dynamics. However, these models also implicitly contain latent information on species and site characteristics. For example, species differences in regional abundance (the row sums) reflect in part differences among species in local resource use (niche differences), a clearly deterministic mechanism in Velland's (2010) scheme and one of the four metacommunity mechanisms apart from mass effects recognized by Shmida and Wilson (1985) and Leibold et al. (2004). Likewise, differences in site richness (or total abundance) must often reflect differences in site conditions. The models thus incorporate mass effects reflecting differences among both species and sites. The site differences (based on column sums) implicitly incorporate effects related to differences among sites in environmental conditions and site proximity or history that affect community size. The "black box" nature of our models means they have the capacity to (implicitly) include various kinds of information on the factors likely affecting community assembly without having to explicitly include any information on them or make assumptions about which are the most important. This approach yields a pair of potentially powerful models based only on the empirical information about the distributions (or abundance) of species over sites. The models do, however, assume independence between species and site effects, excluding all non-additive interactions between whatever factors affect the row versus column sums.

Research to date has not distinguished species mass effects from site mass effects to assess their relative importance or how they might interact. It might be possible to do this by modifying the models presented here. For example, one might construct a species mass effect model based only the variation in species incidence values (or abundance) while excluding column totals from the model (and thus the effects of site richness or total plant abundance).Such a model would test the idea that any species that is present (or abundant) at a site above its meatacommunity mean incidence (or abundance) would tend to disappear (decrease in abundance) at that site while those missing (or less abundant) should colonize (increase in abundance). The accuracy of the predictions of this species effect model could then be compared to those from the two-way model we present. Likewise, one could construct a site effect model based only on variation in species richness (or total plant abundance) among sites. Both one-way models would tend to homogenize species incidence / abundance over species (or site richness / abundance over sites) but might provide a way to assess the relative size of species vs. site effects. It might also be of interest to analyze variation in how individual site x species cell values depart from predicted values. These deviations could, for example, be summed up across rows or columns, allowing us to compare the extent to which individual species (or sites) depart from overall expectations to species' functional traits (or local site conditions). This could generate insights into which mechanics may be driving the dynamics we observe.

Assessing the relative roles of site and landscape conditions, species traits, and stochastic forces on regional metacommunity processes remains a major challenge in community ecology. Actual patterns of colonization, local extinction, and local shifts in abundance surely represent a complex interplay between stochastic and deterministic forces acting at various scales (Germain et al. 2013). Untangling these has proved to be difficult. Simple models like those presented here are useful in that they provide a baseline and standard for judging other, more complex models that incorporate data on species and/or site characteristics. These more complex models are valuable in allowing us to explore how particular assumptions and these ecological details can affect outcomes. We encourage comparing predictions from the models presented here to those with more realistic details on: a) differences among species in niche characters and functional traits; b) differences among species in dispersal ability; c) differences among sites in local environmental conditions, and/or c) differences among sites in landscape conditions or spatial proximity. Like neutral theory (Bell 1991, Hubbell 2001) and other null models (Gotelli and Graves 1996, Hausdorf and Hennig 2007, Stark et al. 2006, Ulrich 2007, Ulrich and Gotelli 2010), the models developed here provide a standard for comparison. To be worth the extra data and effort that more detailed models demand, they should significantly outperform simpler models like the incidence and abundance models presented here.

Ecologists and conservation biologists seeking to exploit metacommunity patterns to predict species turnover and threats to particular species or populations have often been frustrated. Initial hopes that patterns of metacommunity nestedness would serve to predict colonizations and extinctions met with only limited success (Azeria *et al*. 2006; Donlan et al. 2005; Azeria and Kolasa 2008). Our models do not rely on nestedness and, in fact, performed well in two metacommunities lacking significant nestedness (Mudrak 2010). After their exhaustive but frustrating effort to apply six theories to three classes of data on bird and reptile distributions at hundreds of sites in Australia, Driscoll and Lindenmayer (2009) stated that "metacommunity ideas cannot yet be used predictively in a management context." Similarly, in studying changes among 86 southern English woodlands over a 70-year interval, Keith et al. (2011) found metacommunity structure to be stable (like ours) despite declines in beta diversity, concluding that "metacommunity structure [is] not a good landscape-scale indicator for conservation status." We find these conclusions premature and urge others to apply the models developed here before dismissing metacommunity approaches in general. Because species groups respond differently to differences in site / landscape conditions and competitive interactions, we should expect that general patterns will be hard to find.

The models presented are Markovian, but were only tested through a single time interval. We can, in turn, use the 2000s metacommunity matrices to predict the changes expected over the next several decades. Data sets continuing two successive intervals would allow us to compare model accuracy over two intervals of change within a single metacommunity. With stationary transition probabilities, most Markov models converge after multiple intervals to a stable state (e.g., the stable age distribution in Leslie matrix models of population growth). However, we expect stochastic processes acting within sites to regenerate local variability and reset community dynamics before the metacommunity reaches any global equilibrium. Predictions at each time step could still prove valid even though no overall metacommunity convergence occurs. These stochastic forces include storms and other disturbances, local demographic stochasticity, and continual invasions, epidemics, and patchy variation in predation.

Predictions from these models should be tested for their generality in other contexts where extensive repeated survey data exist. If predictions from these models prove to be reliable, ecologists will have gained a useful tool for predicting community change, improving our ability to test hypotheses about the forces driving community assembly. We can test our understanding further using these models as a baseline to see whether and how including additional information on species traits, site conditions, and landscape context in more complex models improves their ability to predict changes in species incidence or community composition. If these models prove to be accurate, conservation biologists could use them as simple tools to predict risks of local species declines or extinctions for particular sites even when they lack data on species and site characteristics. Alternatively, they might conclude that it is best to expend their limited resources to protect sites or recover populations at those sites predicted to be more viable in terms of supporting healthy future populations. It may also prove interesting to compare species in terms of their sensitivity to mass effects inferred from the row or column totals and to compare the accuracy of predictions from these models among landscapes that differ conspicuously in disturbance dynamics or habitat fragmentation.

## Acknowledgements

Our field studies were supported by both NSF (awards DEB-9974041, DEB-0236333, and DEB-1046355) and the USDA Biology of Weedy and Invasive Species program (National Research Initiative award #2008-35320-18680). E.M. was supported by an NSF Graduate Research Fellowship. N. Gotelli, R. Holt, Y. Mikalakis, M.N. Dawson, and anonymous reviewers provided useful suggestions on earlier versions of this manuscript. DMW thanks the ISEM unit of the University of Montpellier, France and the LabEX program for supporting sabbatical time on this project.

